# Logistics of Bone Mineralization in the Chick Embryo Studied by 3D Cryo FIB-SEM Imaging

**DOI:** 10.1101/2023.02.09.527853

**Authors:** Emeline Raguin, Richard Weinkamer, Clemens Schmitt, Luca Curcuraci, Peter Fratzl

**Affiliations:** Department of Biomaterials, Max Planck Institute of Colloids and Interfaces, Am Mühlenberg 1, Potsdam 14476, Germany

**Keywords:** Mineralization, Bone development, Vesicles, Mineral transport, Cryo FIB-SEM, Chick embryo

## Abstract

During skeletal development, bone growth and mineralization require transport of substantial amounts of calcium, while maintaining very low concentration. How an organism overcomes this major logistical challenge remains mostly unexplained. To shed some light on the dynamics of this process, we use cryogenic Focused Ion Beam-Scanning Electron Microscopy (cryo-FIB/SEM) to image forming bone tissue at day 13 of a chick embryo femur. We visualize both cells and matrix in 3D and observe calcium-rich intracellular vesicular structures. Counting the number of these vesicles per unit volume and measuring their calcium content based on the electron back-scattering signal, we are able to estimate the intracellular velocity at which these vesicles need to travel to transport all the calcium required for the mineral deposited in one day within the collagenous tissue. We estimate this velocity at 0.27 μm/s, which is too large for a diffusion process and rather suggests active transport through the cellular network. We conclude that calcium logistics is hierarchical and based on several transport mechanisms: first through the vasculature using calcium-binding proteins and the blood flow, then active transport over tens of micrometers through the network of osteoblasts and osteocytes and, finally, diffusive transport over the last one or two microns.

## 1. Introduction

Bone formation is a complex and well-orchestrated process in which minerals are incorporated into a type I collagen matrix with a multiscale architecture. This arrangement into a nanocomposite constitute the lowest level of a structural hierarchy, from the nano- to the macro-scale, in order to achieve bone’s mechanical, biological and chemical functions [1]. Thus, bone matrix formation and mineralization involves a number of cellular and molecular mechanisms [2]. During bone development, a major challenge is the necessity of transporting massive amounts of ions from the blood stream to sites of mineralization, a transport that has to be strongly controlled due to the necessity of keeping calcium ions in serum and inside cells in the order of millimolar concentrations [3].

The mineral found in bone is carbonated hydroxyapatite [4], a crystalline phase of calcium phosphate that is ultimately responsible for the stiffness of bone but also plays a major role in the maintenance of mineral homeostasis. Our focus in this paper is not on the regulation of this biomineralization process, but on the logistical problems that have to be solved to enable the formation of mineral crystals in sufficient quantities to allow bone growth. In order to elucidate the transport logistics involved in biomineralization, it will be essential (i) to identify the transport path of calcium and phosphate ions from the bloodstream to the mineralization site, (ii) to determine the site at which mineralization initiates and (iii) to describe the form in which the transport of ions occur. The formation of the mineral crystals reportedly follows a precursor phase of amorphous calcium phosphate and is mediated by cells [5], although many details remain to be elucidated [6]. From an historical perspective, an early concept on bone mineralization involves the active transport of extracellular matrix vesicles containing calcium and phosphate ions, whose concentration is sufficient to begin the initial phase of mineralization that will eventually form hydroxyapatite crystals within the vesicles. These crystals will then rupture the membrane due to their continuing growth and be release into the extracellular environment [7]. Later, it has been suggested that mineralization was guided by the collagen matrix, where an accumulation followed by a precipitation of calcium and phosphate ions subsequently forms mineral crystals in the gap regions within the collagen fibrils [8]. More recently, it has been proposed that mineralization results from amorphous mineral precursors transiently formed and deposited within the gap regions of the collagen fibrils and subsequently crystallized into hydroxyapatite [5c]. However, recent studies have challenged some of these mechanisms by showing that mineral ions and precursors are present in the extrafibrillar spaces, where they form aggregates of disordered crystals and also penetrate into the intrafibrillar gap zones [9]. Regardless of the crystallization pathways, the presence of amorphous mineral as cargo within vesicles were found in osteoblasts and bone lining cells in zebrafish [5c, 10], embryonic mouse [11] and in chick embryo [12] using 2D cryo-SEM techniques. The intracellular transport of amorphous mineral within vesicles could be more efficient due to its isotropic and more malleable shape [13]. This mode of transportation through intracellular vesicles and their role in the biomineralization process is better established in some marine organisms such as the sea urchins [14]. However, in the bone of vertebrates, the transport path needed for mineral precursors to find their way to the mineralizing structure across relatively huge distance remains unclear. An efficient transport strategy of mineralization precursors is particularly important during growth and development, where bone is rapidly forming and mineralization needs to increase at a dramatic rate to ensure the mechanical performance of the bone tissue.

The growing chick embryo is a good model to study this bone mineralization logistics for several reasons. First, bone formation is rapid, with earliest sign of bone starting at 8 days of incubation [15]. Second, long bones are small, which allows to image a larger fraction of the bone comprising both the mineralized bone matrix and the cellular environment at high resolution. Third, long bone ossification in birds occurs only through intramembranous ossification: the initial cartilaginous matrix is resorbed instead of being mineralized before remodeling into bone, as it is observed in mammals [16], where bone is largely formed through endochondral ossification from this mineralized cartilage template [17]. Only diametric growth of bone is achieved by intramembranous ossification in both cases [18]. Having only one type of mineralized tissue in the forming skeleton of birds avoids interpretation issues during imaging.

A simple calculation based on this animal model, demonstrates the scale of the logistics problem that has to be solved to mineralize the skeleton (Figure 1). The total amount of calcium ions found in the body of a 2 kg chicken (with a skeleton mass of approximately 0.380 kg) is about 68 g [19] or 1.7 mol. The blood serum, which is the main source of ions transported from the egg shell, has a calcium concentration of 6.75 * 10^−3^ mol/l [20]. Consequently, 252 liters of serum would be required to supply the amount of calcium needed to mineralize the skeleton. The box in Figure 1 represents this substantial volume, which is approximately 1250 times the chicken’s skeleton volume, based on a bone density of 1900 kg/m^3^ [21]. Thus, the mineralization process raises a considerable challenge in terms of throughput of serum and transport of ions from the serum to the site of deposition. While calcium-binding proteins, most notably Fetuin-A [22], are known to support the vascular transport of calcium, the situation is much less clear when it comes to the transport beyond the vascular system, through newly formed collagenous osteoid.

**Figure 1:**
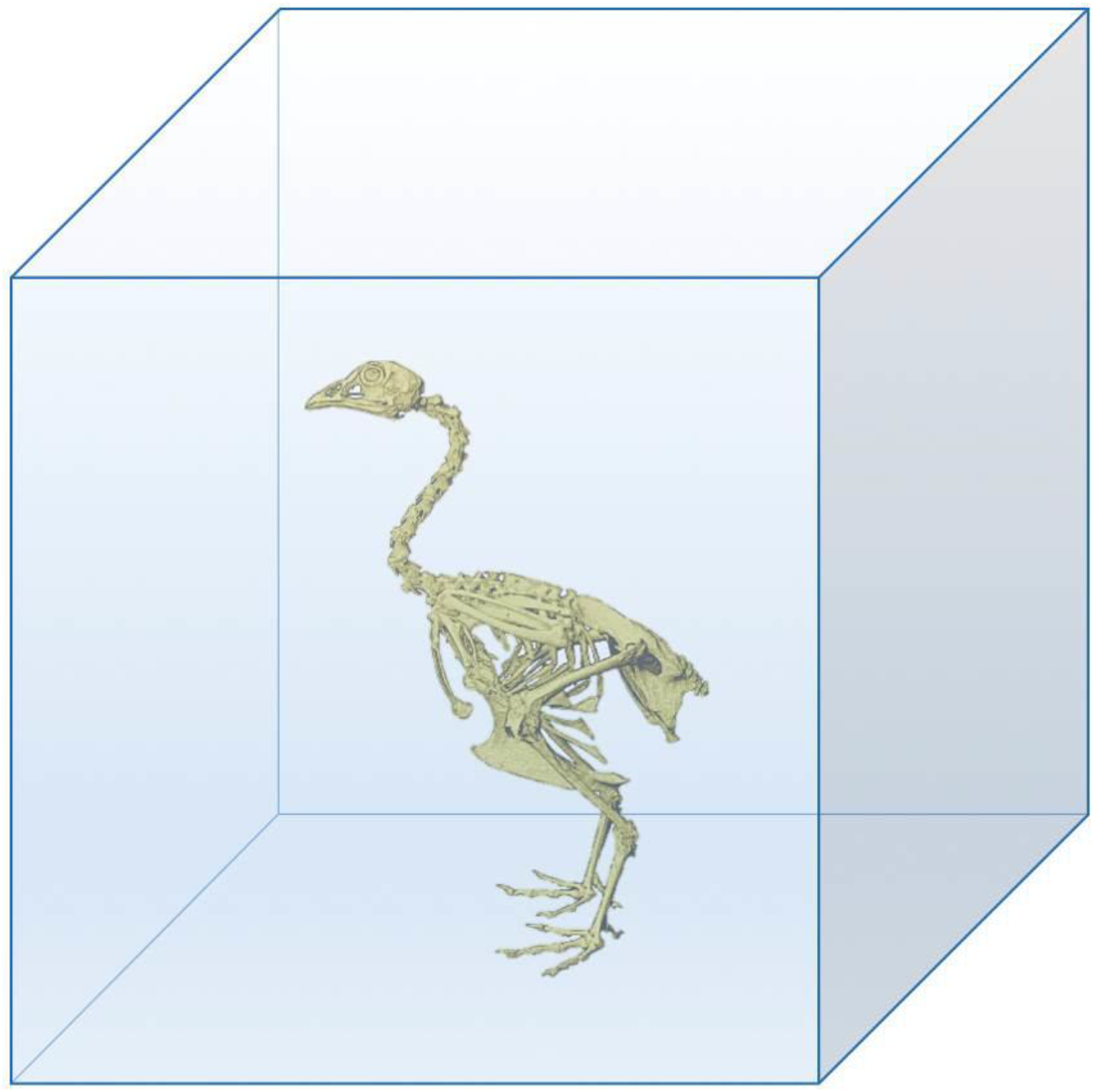
Schematic representation of the volume of blood serum needed (blue box) to provide sufficient calcium to mineralize the skeleton of a chicken. The needed volume corresponds to 1250 times the skeletal volume of the chicken.

In the chick embryo, mobile calcium ions originate from the eggshell and are transported by the highly vascularized chorioallantoic membrane [23]. To ensure the proper signaling function enabled by the calcium, cells need to maintain a very low concentration of calcium in their cytosol, around 100 nM [24], which is even several orders of magnitude less than the concentration found extracellularly. To sustain this low cytosolic concentration, calcium ions are stored in various organelles, such as the endoplasmic reticulum [25], the Golgi apparatus [26] and mitochondria [27], which then release calcium ions through channel systems [28]. In bone cells, the calcium storage is even more critical since large amounts need to be transported to fulfill the mineralization requirement.

A study of the mineralization of forming human osteons demonstrated the involvement of the lacuno-canalicular network in the mineralization process and led to the interpretation that this network might be the source of both mineralization precursors and inhibitors diffusing through the matrix [29]. On an even smaller length scale, it has been proposed that mineralization in turkey leg tendon [9] and in mouse bone [30] is facilitated by a network of nanochannels found between the collagen fibrils (of approximately 40 nm in diameter compared to 300 nm for the diameter of a canaliculus).

While the routes of mineral transport became clearer over the last decade, a much more challenging problem is to image the mineral precursors that are transported. Most likely, the transport involves bone cells, which therefore have to be preserved during sample preparation. In addition, most of the studies on mineral transport were based on 2D images, which makes it hard to determine the spatial distribution and the quantification of the mineralization component [5c, 10–12, 31]. When performed in a 3D volume, the studies used dehydrated samples or freeze substitution preparation that do not allow the preservation of the precursors phase [9, 29].

In this study, we use high-resolution FIB-SEM with the serial surface view method in cryogenic condition to visualize in 3D the mineral precursors, which are transported towards the site of mineralization in the forming bone of the chick embryo. This technique enables the visualization and quantification of mineral precursors, cells, as well as the mineralized bone matrix with samples examined under condition that are close to their native state. A quantification of the obtained “snapshots” allows us to interpret them dynamically, i.e. to estimate the speed at which mineralization precursors have to be transported from the vascular system to the site of mineralization.

## 2. Results

### 2.1 Characterization of the chick embryo femoral microarchitecture

To evaluate the increase of mineralized bone volume during the time interval of one day, chick embryos at 13 and 14 days post-fertilization (known as embryonic developmental day (EDD)) were imaged using microcomputed tomography (micro-CT) (Figure 2A and B). For each femur, a region of interest 1 mm thick, centered at the midshaft (Figure 2A-B red and orange rectangle) was analyzed to account only for the intramembranous bone apposition. The microscopic structure of the bone is characterize by bony plate-like elements spreading radially from a well-defined bone collar that surrounds the marrow cavity, creating a three-dimensional trabecular network. The appositional bone growth is asymmetric and occurs predominantly in the posterolateral region in these two embryogenic stages. An increase of 73% of the mineralized bone volume was observed between EDD13 and EDD14, from 0.155 mm^3^ to 0.268 mm^3^ respectively. Backscattered (BSE) image of the cross-sectional midshaft at EDD13 (Figure 2C) shows that the innermost bone collar is brighter, e.g. more mineralized while the mineral density decrease towards the periosteal side especially in the preferential growth direction. Numerous osteocyte lacunae can be observed inside the bone collar and struts. The hematoxylin and eosin (H&E) staining (Figure 2D- E) reveals a periosteal layer rich of osteogenic cells. Remnants of primary cartilage in the marrow cavity are still present at EDD13 since bone develops on the surface of the cartilage in birds. This cartilage does not mineralize and is never invaded by blood vessels (Figure 2D, black arrows) [16]. In the spaces in-between trabeculae, elongated cells form blood vessel lumina (Figure 2E), as previously reported [12].

**Figure 2:**
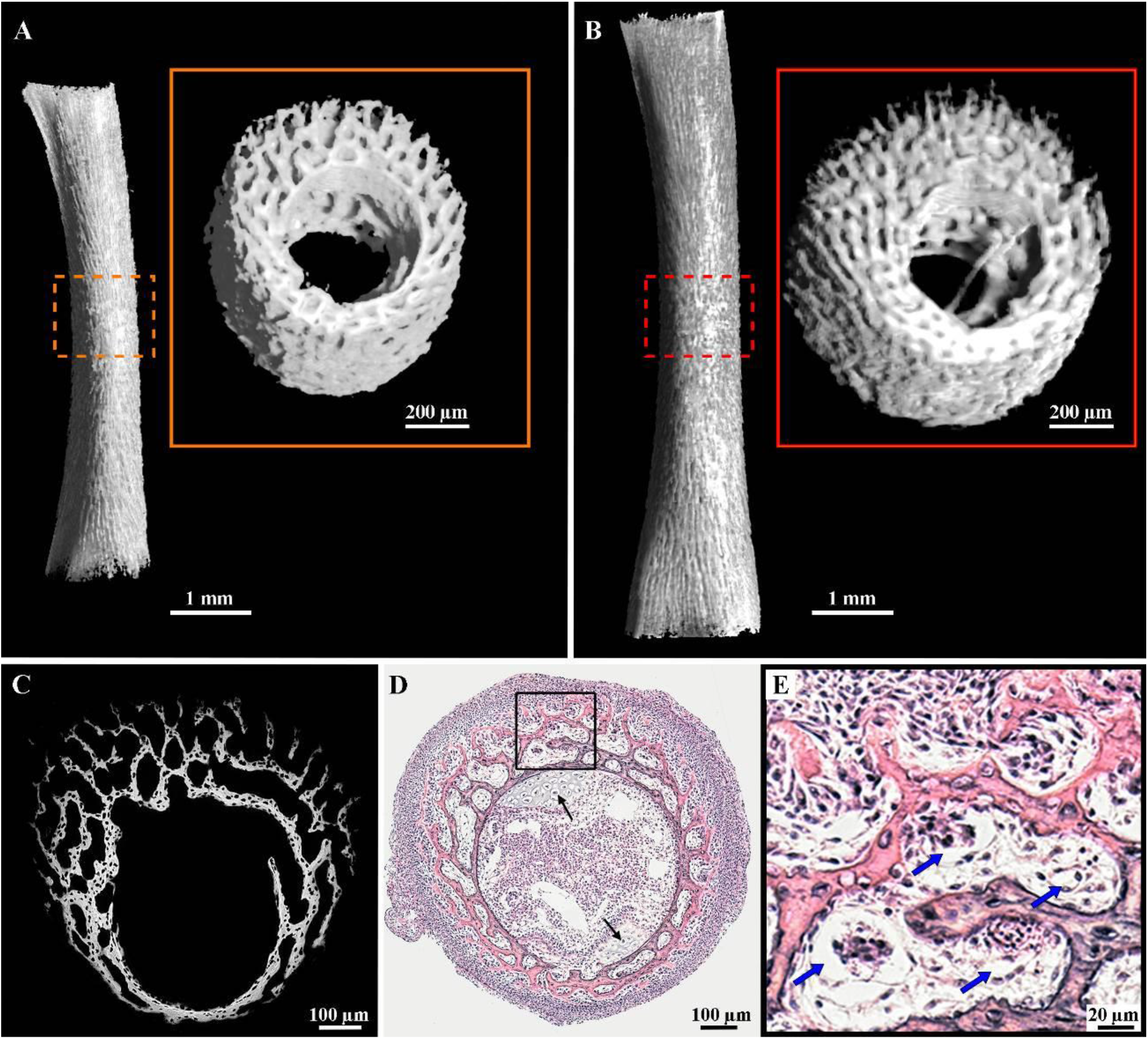
Morphology of the chick embryo femur at the microscopic level. (A-B) 3D orthographic projection in the anterior view of the mineralized right femoral bone at EDD13 (A) and EDD14 (B) obtained by micro-CT. The orange and the red dotted box in each image highlight the region of interest of identical dimensions from which the mineralized volume was computed. These ROIs are shown in higher magnification in the orange and red box. (C) Representative BSE image of the midshaft cross-section at EDD13 showing variations in the grey scale intensities. Brighter regions depict more mineralized areas while darker regions, located on the outermost area where the new bone formation occurs (periosteal side), are less mineralized. (D) Representative hematoxylin and eosin-stained femoral midshaft cross-section at EDD13. Remnants of cartilage persist in the marrow cavity, inside the bone collar (black arrows). (E) Higher magnification of the area delimited by the black square in (D) showing an abundance of osteogenic cells at the periosteum (stained in light purple with the nuclei in dark purple) and newly formed, less mineralized, trabeculae (light pink). The blue arrows indicate blood vessels lumina.

### 2.2 Presence of intracellular vesicles containing mineral precursors in cells associated with partially mineralized bone matrix

To investigate bone mineralization logistics in the rapidly growing chick embryo, the femoral midshaft of three different specimens were imaged using Focused Ion Beam with Scanning Electron Microscopy (FIB-SEM) and the serial surface view (SSV) method in cryo conditions. The region of interest analyzed in all stacks was located at the sites of most active bone growth, namely in the outermost bone trabeculae.

Figure 3 shows bone cells surrounded by mineralized matrix imaged in the electron microscope with two different detectors, which provide complementary information and should be therefore considered together. The mixed Inlens/SE detector allows to visualize the structure of the sample due to a contrast resulting from differences in the local surface potential while the contrast in the BSE detector results from the electron density differences in the material. While one cell is already embedded in the mineralized matrix (Figure 3A) and could therefore be considered an osteocyte, the cell in Figure 3B is still in contact with unmineralized collagen (white arrow). Within the cells, spherical structures, which display a well-defined membrane and are therefore identified as vesicles, can be observed. These vesicles contain granular structures of 80 nm diameter on average (Figure 3A and B, arrowheads), which correspond to electron-dense particles when seen with the BSE detector (Figure 3C and D, arrowheads) (see also Movie S1). Interestingly, not all the granules that can be seen in the mixed Inlens/SE images displays this type of brighter contrast in the BSE images.

**Figure 3:**
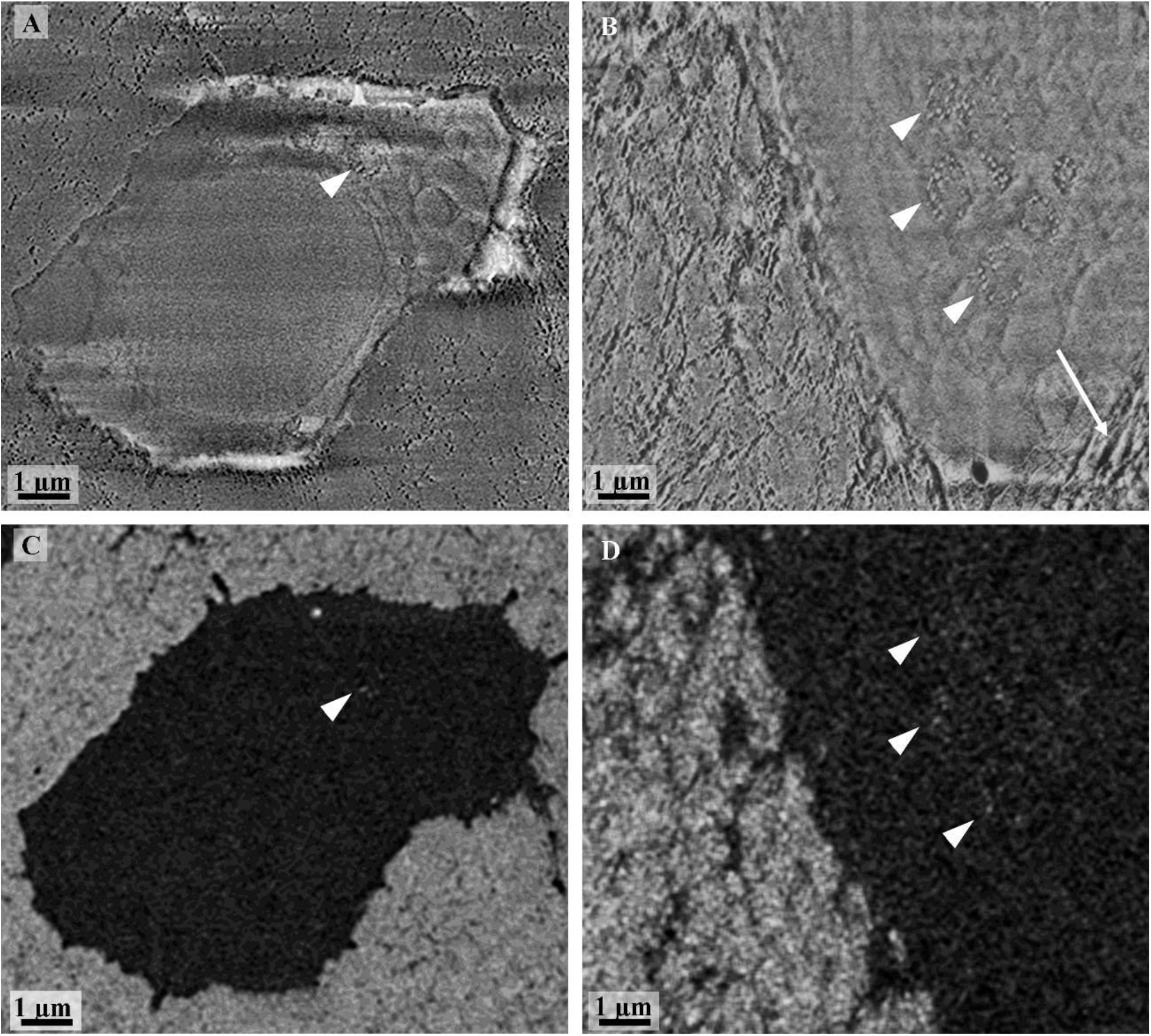
SEM images of the intracellular vesicles containing mineral precursors (arrowheads). The mixed Inlens/SE electron detector shows the presence of granules in (A) an osteocyte surrounded by the mineralized matrix and (B) in a cell at the interface between the mineralized and unmineralized matrix (arrow). The intracellular vesicles are well defined by their membrane. Mineral precursors are found inside some of the vesicles and appear brighter in the corresponding BSE images (arrowhead) (C, D).

In addition, Figure 4 demonstrates that many vesicles do not contain dense particles (marked in yellow), i.e., no granular structures in the mixed Inlens/SE image (Figure 4A) and no dense particles (brighter spots) in the BSE image (Figure 4B). For our evaluation, we only considered the vesicles that contained electron-dense material identified as mineral precursor.

**Figure 4:**
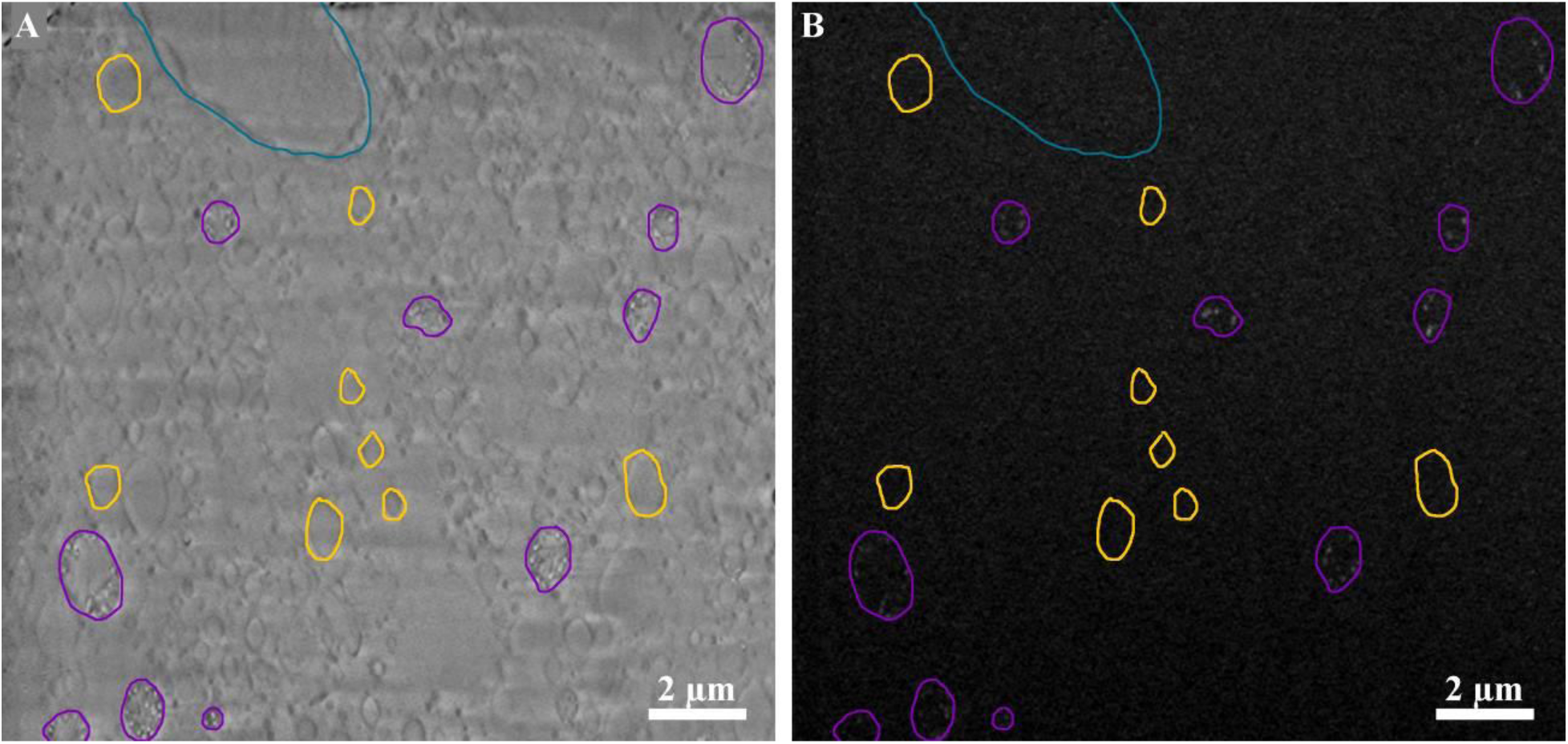
Mixed Inlens/SE (A) and BSE (B) images of stack 2 showing a cell nucleus (circled in turquoise) and the numerous vesicles contained inside a cell. The vesicles containing mineral precursors are circled in purple. Note that these vesicles containing mineral precursors can be distinguished also in the mixed Inlens/SE image (A) by the presence of granules compared to the vesicles that does not show any mineral precursors (some examples are shown in yellow).

3D imaging allows quantifying the volume of the different structures. We investigated three different bone samples for which we obtained one 3D stack each. The mean volume of the vesicles containing minerals (all stacks combined) was 0.65 μm^3^ and, more specifically, it was 0.60 μm^3^ for stack 1, 0.74 μm^3^ for stack 2 and 0.26 μm^3^ for stack 3 (Table 1). However, 80% of the vesicles have a volume lower than 1 μm^3^ in all stacks (Figure 5A). We asked the question whether the volume of particles per vesicle or the filling degree of the vesicles (i.e. the ratio of the vesicle and the mineral precursor volume) is the more controlled parameter. Considering a total of 432 vesicles in all stacks combined, we found that the volume of the vesicles is positively correlated with the volume of the mineral precursors contained inside the vesicles (Figure 5B). Fitting the data with a power law relationship, i.e. with a straight line in a double logarithmic plot (Figure 5B), gave for all three image stacks an exponent very close to 1. This indication of a constant filling factor is confirmed by Figure 5C, which does not show any trend of the filling factor with vesicle size. The filling factor is 9% on average and lies in a range between 8 and 10% between the three stacks (Table 1).

**Table 1:** Quantitative data for all three stacks investigated. All volumes are expressed in μm^3^. The volume of vesicles and the volume of mineral precursors inside the vesicles represent the total volume per stack. *Note that the lower volume imaged in stack 3 as mentioned previously results in an underestimation of vesicles per cell.

|  | Stack 1 | Stack 2 | Stack 3 |
| --- | --- | --- | --- |
| Volume imaged | 10 471.33 | 12 086.04 | 1 563.27 |
| Volume of mineralized bone | 2 734.51 | 2 626.39 | 767.87 |
| Number of cells | 6 | 6 | 7 |
| Number of vesicles | 122 | 270 | 40 |
| Average volume of vesicle | 0.60 | 0.74 | 0.26 |
| Filling factor of vesicle | 0.094 | 0.085 | 0.101 |
| Volume of mineral precursors per vesicle obtained with the filling factor | 0.056 | 0.064 | 0.024* |
| Number of vesicles shed by a single cell (#vesicles/min) | 63.2 | 55.3 | 147.6* |
| Release rate of vesicles (time in s to release a vesicle) | 0.95 | 1.08 | 0.41* |

**Figure 5:**
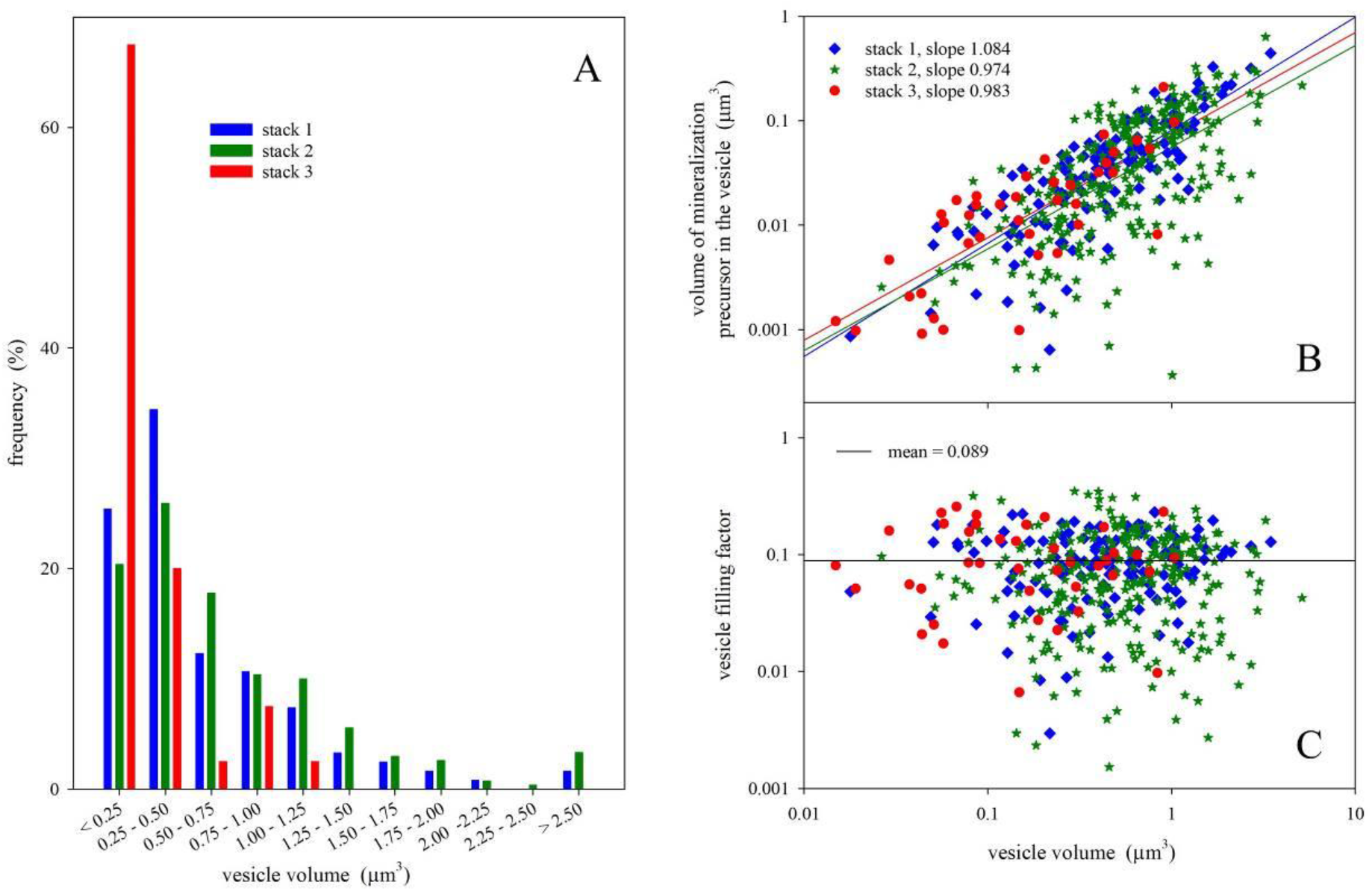
Quantitative data of the vesicles containing minerals precursors. (A) Histogram showing the relative number of vesicle according to their volume for each stack. (B) Relation between the volume of the vesicles and the volume of mineral precursors contained in the vesicles for all stacks presented in a double logarithmic plot. In all three stacks, a fit with a power law equation resulted in exponents very close to 1, which would be expected for a constant filling factor. (C) The value of this filling factor corresponds to 9% on average for the three stacks combined.

Figure 6 shows a 3D perspective rendering resulting from the segmentation of stack 2. A high density of mineral precursor containing vesicles can be observed in the volume imaged and these vesicles are found virtually everywhere around the mineralized bone matrix (Figure 6A). However, they are exclusively located inside cells (Movie S1), and this observation was consistent in all three image stacks; no extracellular vesicles could be observed. Despite the very different (imaged) volume of the cells, the density of vesicles per cell volume was consistent for all three stacks with 0.037 vesicle/μm^3^ on average (Figure S1).

**Figure 6:**
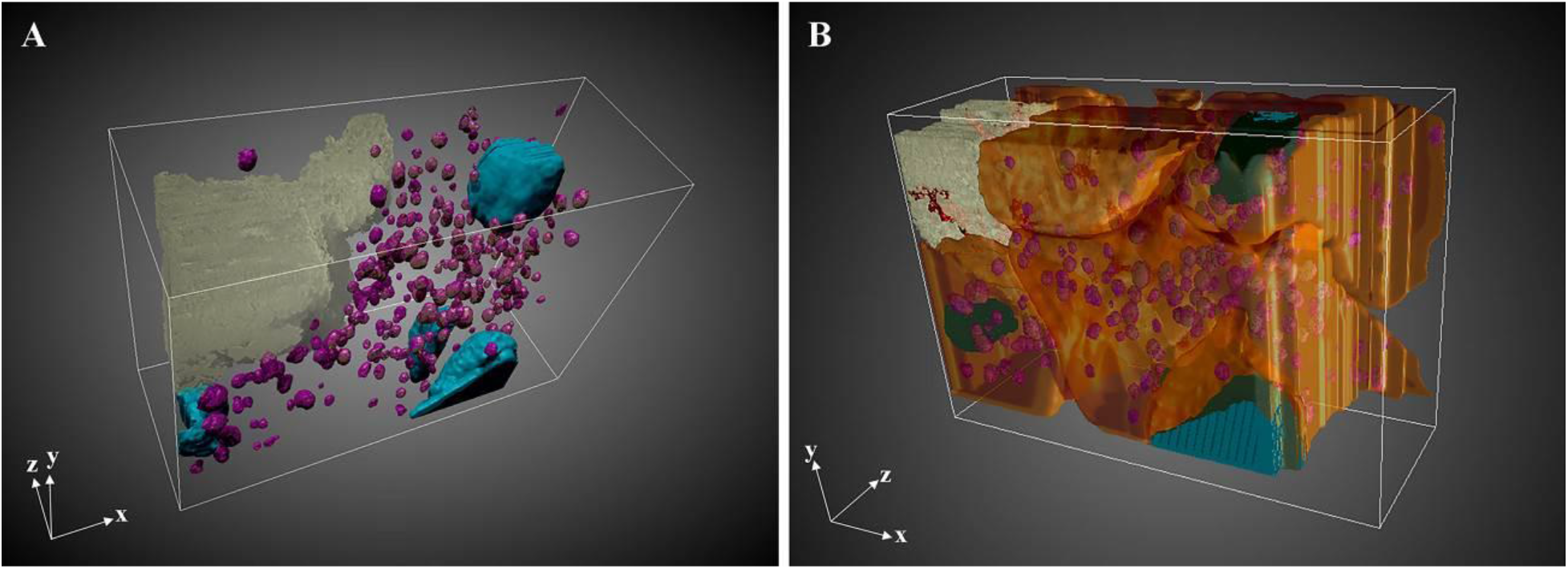
Perspective rendering of the 3D FIB-SEM image stack 2 after manual segmentation of different structural features. (A) Numerous vesicles containing mineral precursors (purple) can be observed near the mineralized bone matrix (light gray yellow) in the volume imaged, delimited by the white bounding box. Nuclei are shown in turquoise as landmarks. (B) The vesicles containing mineral precursors are exclusively found in cells (orange) lining the mineralized bone matrix, at the interphase with the unmineralized matrix. Pre-osteocytes can be observed with canaliculi (red) penetrating the mineralized matrix.

In all three stacks between 6 and 7 cells have been partly imaged, the number of cells being also confirmed by the presence of well-defined nuclei (Figure 6, turquoise). Based on their spatial position, the cells differed in their embeddedness in the mineralized matrix. In accordance with the differentiation from an osteoblast to an osteocyte, cell embedding went along with the formation of the characteristic cytoplasmic extensions, which are accommodated in canaliculi.

### 2.3 Interconnected nanochannels and canaliculi network in the mineralized bone matrix

Figure 7A and B show an example of an image from the bone matrix in the same stack as Figure 6. Since embryonic bone is characterized by a woven structure, defined by a disordered arrangement of the collagen fibrils [32], the 67 nm D-banding pattern is seen only in some areas while other regions appear uniform (Figure 7A). Non-mineralized collagen fibrils are also observed (Figure 7A, black arrowhead). The mixed Inlens/SE image reveals a texture composed of big elongated structures of approximately 300 nm diameter (Figure 7A and B, blue arrows). In addition, smaller channels of 40 nm in diameter are observed in the mineralized bone (Figure 7A and B, white arrows). The BSE image (Figure 7B, white and blue arrows) shows that the contrast of both these structures is dark compared to the surrounding bone material that displays a bright contrast, indicating that this material is less mineralized. When viewed in 3D (Movie S2, Figure 7C and D, red), the bigger structures correspond unequivocally to the canaliculi since they are connected to the cells lining the mineralized matrix. Interestingly, the smaller structures are forming an interconnected meshwork of nanochannels localized between the collagen fibrils (Movie S2, Figure 7C and D, blue) that represents more than 14% of the volume of the mineralized bone. The nanochannels are associated to the canaliculi.

**Figure 7:**
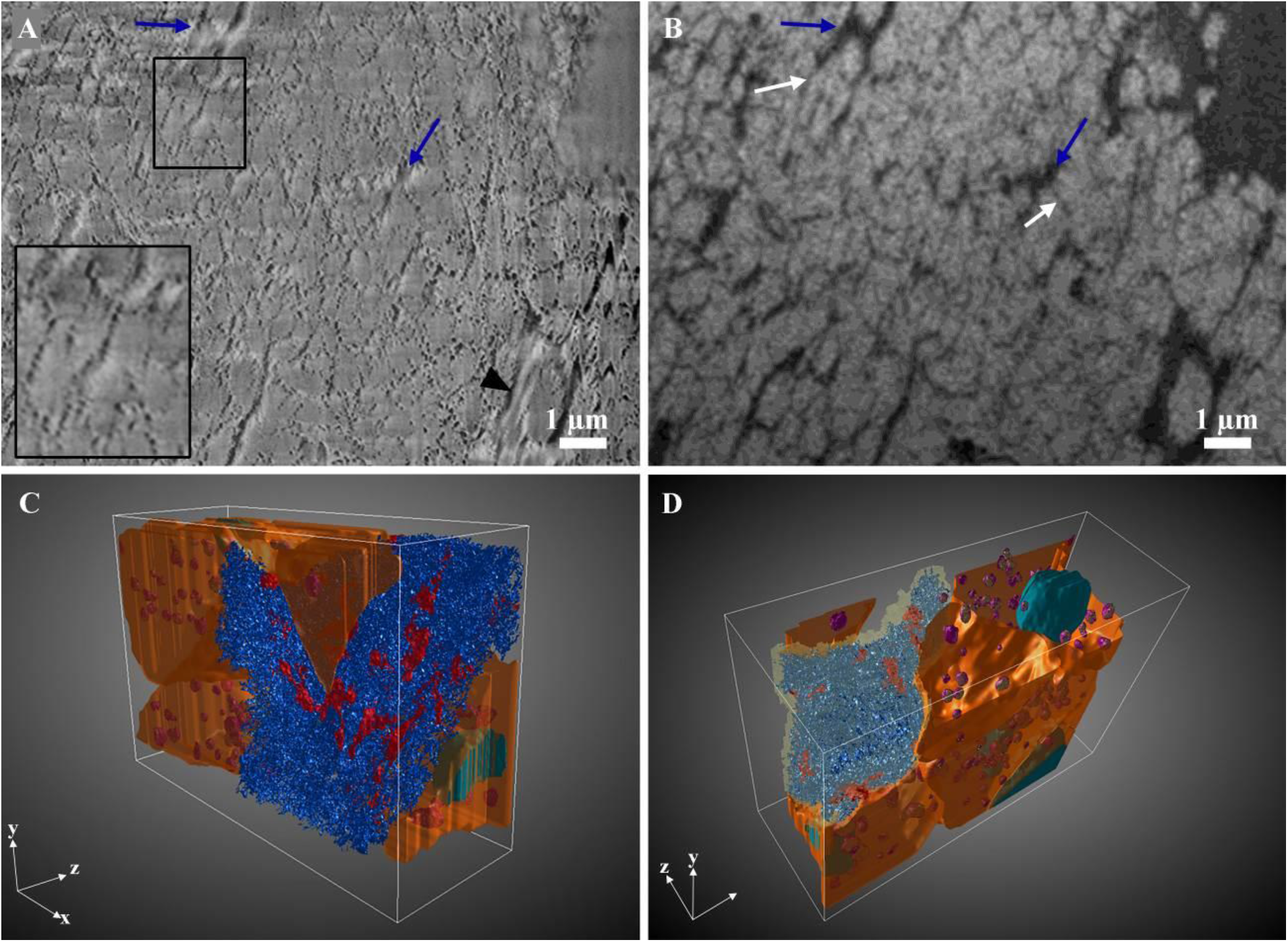
SEM images of the mineralized bone matrix in stack 2. (A) Image of the mixed Inlens/SE electron detector showing canaliculi (blue arrows) as well as nanochannels (white arrows). Non-mineralized collagen fibrils (black arrowhead) and mineralized collagen fibrils (lower left inset) can be seen with the 67nm D-banding. (B) Corresponding BSE image of (A) highlighting that the nanochannels (white arrows) are displaying a darker contrast. These channels branch to the canaliculi. (C) 3D perspective rendering showing the segmentation of the interconnected network of nanochannels (blue) and canaliculi (red) in relation with the cells (orange). (D) The mineralized bone is seen in light yellow. The numerous vesicles containing mineral precursors (purple) can be observed inside the cell and around the nucleus (turquoise).

### 2.4 Estimation of vesicle velocity for the mineralization in the chick embryo femur

To interpret the 3D snapshots dynamically and to estimate kinetic parameters, we used the quantification of the different transport structures observed in each stack from the segmented data. We formulate the problem of mineralization logistics in analogy to an inverse transport problem where trucks are loaded with material. (i) The total amount of transported material within one day is known. (ii) Knowing how much material a single truck can load, the total amount of transported material can be expressed as the number of trucks that are needed to arrive per minute. (iii) A snapshot of the number of trucks on a piece of the transport highway allows to assess how fast the trucks have to move in order to deliver the needed material. Obviously, in this analogy trucks correspond to vesicles, which are imaged in a 3D volume and the loaded material per truck to the amount of mineral precursor per vesicle. In the calculation, we assume that the mass density of the mineral precursor within vesicles is similar to the mass density of the bone mineral phase, so that the calculation can be limited to volumes (instead of masses).

(i) In a first step, the volume of bone to be mineralized per cell in one day has to be calculated. Starting point is the μCT-measurement of the osteocyte lacunar density (Movie S3) of 196 000 lacunae/mm^3^. Thus, the average volume a cell needs to mineralize corresponds to 5100 μm^3^ or to a cube with an edge length of 17.2 μm. Comparison of the μCT at day 13 and 14 reveals that at the outer fringe of the femur, the region imaged with the FIB-SEM, new trabeculae were formed within this one day with an average thickness of 16 μm (Figure 8) that is really close to the estimated edge length of the mineralizing volume. (ii) Combining data from all stacks yields an average volume of a vesicle of 0.65 μm^3^ and a filling factor of 0.089, therefore, the volume of mineral precursor per vesicle is 0.058 μm^3^. The number of vesicles that have to be shed by a single cell per minute to mineralize a volume of about 5100 μm^3^ in one day equals to 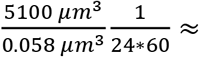 61, i.e. about one every second. (iii) To estimate the average number of vesicles per cell, we have to take into account both the strongly reduced size of a (pre-)osteocyte embedded in collagenous tissue as compared to an osteoblast [33] and the density of vesicles containing precursor in the partial cell volumes imaged in the three stacks (Table S1). For a pre-osteocyte, we assume a volume of 1000 μm^3^ and combining the data from all 19 partly imaged cells, we obtain a vesicle density (#vesicles per μm^3^) of 0.037 μm^−3^ (Figure S1), resulting in 37 vesicles per pre-osteocyte. Assuming the distance that vesicles need to travel from the trabecular surface to the site of mineralization inside the trabecula to be about 10 μm (that is, a bit more than half the thickness of the trabecula), the average velocity of a vesicle within the cell can be estimated as 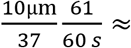 0.27μm/s. The values calculated separately for each stack are presented in table 1.

**Figure 8:**
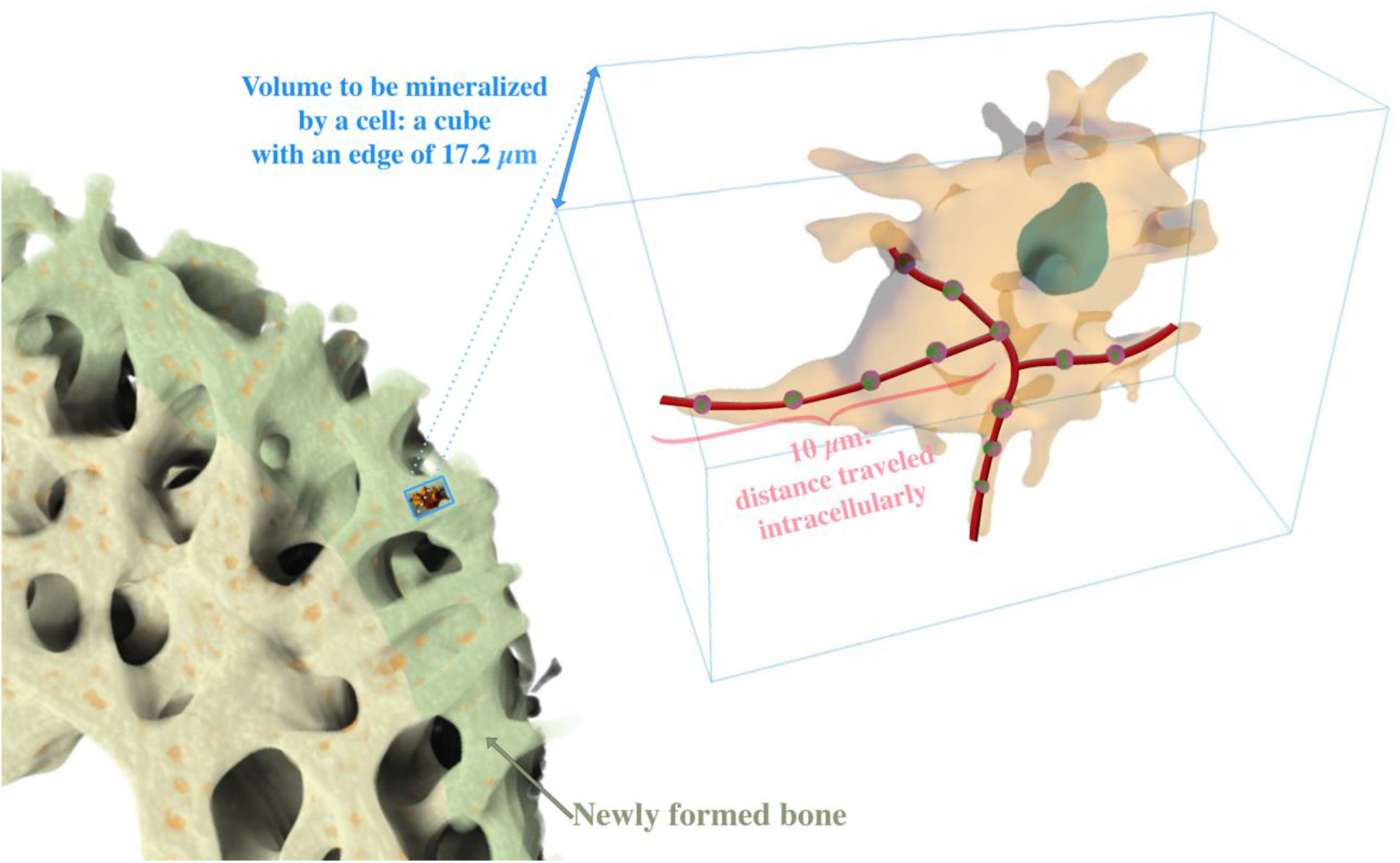
Schematic illustration of the volume that has to be mineralized by a cell. The intracellular velocity of the vesicles (pink-green spheres) is obtained from their number within a cell, an assumed intracellular distance travelled of 10 μm (traveling path in red) and a calculated shedding rate of vesicles per minute (see text).

## 3. Discussion

The rapid growth of bone in developing avian and mammalian organisms involves the task to transport substantial amounts of calcium from the blood stream to the sites of mineralization. 3D imaging using FIB-SEM under cryo condition provides a new view into the details of this transport process. With the chick embryo femur as model system, we could not only show the presence of vesicles containing mineral precursors inside the bone forming cells and osteocytes in their close to native state in 3D. In a painstaking analysis of the images evaluating hundreds of vesicles, a statistically reliable estimate was obtained of how much mineral precursors a single vesicle is transporting on average. Relating this value with the total amount of calcium needed for the mineralization during a day of embryonic growth (specifically, between embryonic day 13 and 14) and evaluating the number of bone cells actively involved allows to assess the number of vesicles that has to be shed per hour to ensure a sufficient calcium transport: 61 vesicles per minutes approximately.

The presence of vesicles containing mineral precursors inside the cells juxtaposing the mineralized bone matrix has been observed in different animal models using cryo-SEM methods that do not required chemical treatment [5c, 10–12]. To our knowledge, it is the first time that they are reported in 3D and in a large volume, allowing both a spatial visualization and a quantification of their occurrence. The size of the precursors-containing vesicles were found to be much more variable than previously reported, ranging from 0.3 to 2.8 μm in diameter, but are still consistent with studies in the chick embryo [12a]. In both the zebrafish caudal fin bone and the mouse calvaria, intracellular vesicles containing mineral are reported with a diameter of 1 μm or less [10–11]. While the average diameter found here was similar, the higher density of vesicles observed in our study might explain the differences but we cannot exclude variation among vertebrates.

The question arises whether all the observed vesicles are actually involved in the mineralization process. In an in vitro study, Boonrungsiman et al. [34] established a direct transportation of calcium ions and potentially phosphate ions between intracellular vesicles and mitochondria in cultured osteoblasts, suggesting a transfer mechanism by diffusion between the two entities. Still based on cell culture, other researchers found that the transport between intracellular vesicles and mitochondria involves the role of lysosomes, where the latter can fuse and degrade the membrane of the mitochondria, freeing the mineral ions while preventing crystallization due to their acidic nature. Lysosomes can thus act as intracellular transporter for mineralization and release their content by exocytosis [35]. The endoplasmic reticulum is well known to facilitate intracellular trafficking and is the main reservoir of calcium ions in the cell [25]. Through contact sites with the Golgi apparatus, mitochondria and lysosomes, the endoplasmic reticulum can export the calcium ions but also generate myriad of vesicles responsible for distributing their protein content [36]. Here, we show that numerous vesicle both with and without denser particles were found inside the cells. However, we only look and quantify the vesicles displaying a high electron density (denser particle with higher atomic content) from the BSE signals and given the low concentration needed in the cytosol, we are confident that these vesicles are responsible for the mineralization of bone, although we cannot exclude an overestimation. We established that the electron-dense mineral precursors contained inside the vesicle occupied less than 10% of the volume. Various enzymes and proteins acting as inhibitors of crystal formation, growth and calcium binding have been found in the matrix of vesicles [37] that might contribute to ease the releasing of the content.

In addition, we observed only intracellular vesicles but extracellular vesicles containing mineral precursors have been found in calcifying cartilage [7b, 38], dentin [39] and turkey leg tendon [9a, 40] using conventional preparation techniques. The diameter of the vesicles, when reported, was found to be much smaller ranging in overall between 0.09 and 0.2 μm. Thus, it was proposed that these extracellular vesicles were released through cell processes [9a, 39–40]. The size of the vesicles reported here would rather suggest that the mineral precursors-bearing vesicles cannot transit directly through the cell processes of (pre-)osteocytes in bone. Alternatively, it might be also conceivable that the intracellular vesicles reported here contain precursors in a condensed liquid phase, akin to the polymer-induced liquid precursor (PILP) process used in bioengineering [41] and referred usually as the liquid-liquid phase separation (LLPS) theory [42]. This model propose that a mixed fluid could separate into a dilute phase liquid and a dense phase, the latter being composed of amorphous clusters, in accordance with previous studies showing the presence of amorphous calcium phosphate inside the vesicles [5c, 10, 12, 31]. Although this has yet to been shown in bone, this liquid phase has been reported in the biomineralization process from calcium carbonate in the sea urchin [43]. Such mechanism could explain how intracellular vesicles containing mineral precursors of various sizes could transit through cell processes, the liquid phase allowing large deformation of the vesicles.

Although we did not visualize any vesicles with mineral precursors inside cell processes, these processes offer intercellular routes of transport since osteoblasts/pre-osteocytes and osteocytes are connected. Assuming that these vesicles have to be transported across the cell before they and their content are shed into the surrounding matrix, we have calculated the intracellular velocity of the vesicles to be approximately 0.3 μm/s. Such a high velocity cannot be achieved by passive diffusion, and thus, requires an active process during bone formation [44]. In vitro studies showed velocities in the same order of magnitude for cargos enclosed in a fluid membrane, between 0.6 and 0.8 μm/s [45]. Our results are also in agreement with the velocity of molecular motors involved in vesicle transport (0.2 μm/s) [33b].

In order to reach mineralization sites that are further from the lacunae and canaliculi, secondary routes of transport for the mineralization must exist. We observed in the chick embryo femur, a network formed in the mineralized bone matrix of nanochannels. The nanochannels found here appears to be a good path to diffuse the mineral content, whether in the form of ions or precursors. However, their diameter is definitely too small to transport vesicles. Therefore, we suggest that once the vesicles leave the cell, they shed their content in the surrounding matrix. Recent studies found these nanochannels in the growth plate of the mouse femur [30], in human osteonal bone [46] as well as in the turkey leg tendon [9a]. In the turkey tendon, these nanochannels were observed in close relationship with possible matrix vesicle, suggesting that these nanochannels play a critical role in the transport of mineral ions and/or mineral precursors. In the current study, we were able to image both the mineral-bearing vesicles and this network. It has been speculated that these nanochannels were filled with organic content [30, 46]. Similarly, it has been concluded from mineralization patterns that the canaliculi should be sources of inhibitors [29]. Given the average diameter of the nanochannels (around 40 nm) could allow the transport of calcium and phosphate ions while preventing the mineral to precipitate, while the canaliculi could facilitate the transit of mineral precursors.

We proposed a hierarchical model for the mineralization of forming bone based on active transport of vesicles intracellularly (Figure 9). These vesicles might originate from the cell organelles such as the endoplasmic reticulum where they can filled/refilled their cargo. These transporters containing mineral precursors are trafficked intracellularly at a high velocity, and can be shed in the bone matrix through exocytosis where their content subsequently passively diffuse through the nanochannel network. This allows the necessary efficient transportation over substantial distances to reach the mineralization sites. Since the estimated intracellular velocities of the vesicles needed for mineralization are in a range feasible for cells (0.27 μm/s), we conclude that a transport of mineral precursors exclusively via vesicles is sufficient to perform the mineralization task if the bone cells unleash their full transport capacities.

**Figure 9:**
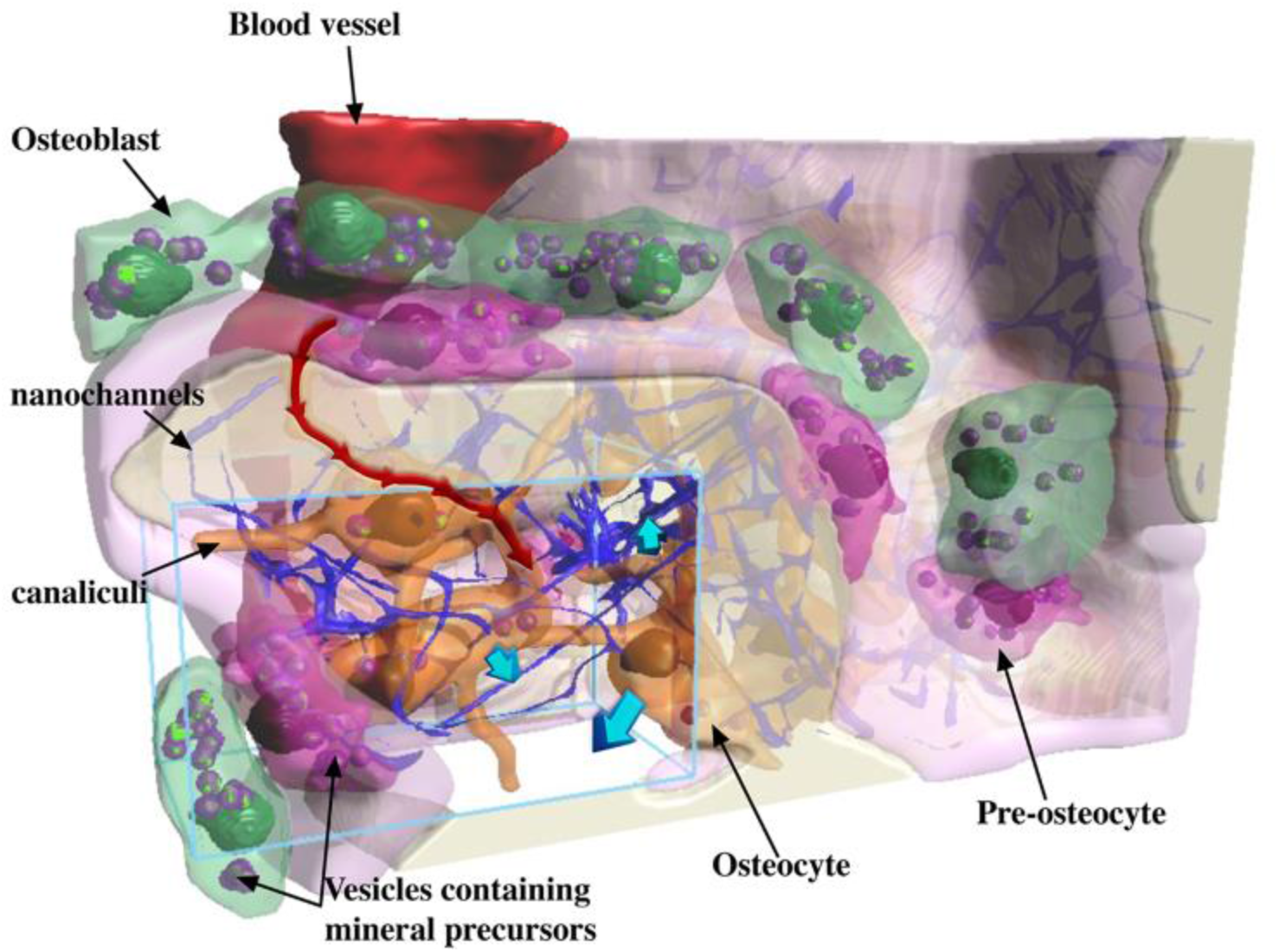
Scheme of the proposed transport model of mineralization in forming bone. The blue rectangular volume shows approximately the area that was investigated using the FIB-SEM in cryo condition, the light pink layer represents the osteoid and the mineralized bone is shown is white. The cells contain numerous vesicles comprising mineral precursors. These vesicles are transported at high speed within the cell bodies and processes (red arrows). A passive transport process might occur when vesicles shed their content into the surrounding matrix and ions/mineral precursor diffuse through the nanochannels network in order to reach the sites of mineralization. The flux of mineral precursors from the cells to the surrounding matrix (represented by blue arrows) was assessed based on a quantification of the 3D image dataset.

We want to emphasize that the performed calculations can only provide an approximate result since they are based on assumptions and use quantitative input from a limited image volume. It is possible that not all the vesicles containing mineral precursors are directly involved in bone mineralization, but some could be a reservoir to maintain a low cytosolic calcium concentration. Therefore, we would overestimate the number of vesicle involved in bone mineralization and the remaining ones would need to move even faster. We are surprised by the low filling factor of vesicles by mineral precursor. Here it is possible that with our methods we detect only the electron-densest part of the precursor and the vesicle contains more precursor in a more diluted form. This underestimation of the vesicle cargo would result in a slowing down of the vesicles. Such a compensation of over- and underestimations make us confident that our calculation provide at least the correct order of magnitude of how many vesicles a bone cell has to shed per minute to enable mineralization and how fast they have to move within the cell.

## 4. Experimental Section

### Eggs procurement and incubation

In accordance with the German Animal Welfare Act and the Laboratory Animal Welfare Ordinance, no approval by an ethics committee for animal experimentation was required. Fertilized Lohmann Selected Leghorn (LSL) chicken eggs (*Gallus gallus domesticus)* were purchased from a commercial hatchery (LSL Rhein-Main Geflügelvermehrungsbetrieb, Dieburg, Germany). Upon delivery, eggshells were gently cleaned with a 70% ethanol wipe to avoid microbial infection, placed in an automatic digital egg incubator (Ovation 56 eco egg incubator, Brinsea, North Somerset, UK) at 37.5 ± 0.5 °C and 55 ± 5% relative humidity, and turned over every hour. Embryos were sacrificed by cervical dislocation and femurs were surgically removed at stage 39 of development according to Hamburger and Hamilton (HH) system [47] (embryonic developmental day EDD13 post fertilization). In addition, one chick embryo was incubated until the stage HH40 (EDD14) to assess the increase of mineralized bone volume in 24h by micro-CT.

### Microcomputed tomography (micro-CT)

One femur at EDD13 and at EDD14 were placed in plastic tube filled with 70% ethanol and imaged with a micro-CT EasyTom 150/160 system (RX Solutions, Chavanod, France). The micro-focus tube was combined with a CCD camera and the X-ray source was set to a current of 250 μA and a voltage of 40 kV and the voxel size ranged between 4 and 7 μm. The 3D reconstruction from the two-dimensional projections images was performed using the X-Act software (RX Solutions, Chavanod, France). The mineralized bone volume was calculated on a region of interest of 1 mm thick centered in the midshaft for each femur from a threshold-based segmentation. The threshold to separate the mineralized vs. non-mineralized phase in each image of the ROI has been found by estimating the point of a horizontal slope in the gray level histogram of each image slice based on calculating the second derivative. The global threshold is then defined as the mean position of these points in all images, and their standard deviation is used to quantify the uncertainty of the threshold. The volume of the bone was computed from the threshold-based segmentation in Dragonfly software according to the previously established threshold (Object Research Systems (ORS) Inc, Montreal, Canada).

To compute the density of the osteocytes lacunae, one femur at EDD13 was fixed and contrast-stained with phosphotungstic acid (PTA) (Aldrich chemicals, 12501234) following the protocol developed by Metscher [48]. The sample was imaged using the nano-focus tube available in the micro-CT EasyTom 150/160 system (RX Solutions, Chavanod, France), combined with a flat panel at a current of 82 μA, a voltage of 80 kV and with a voxel size of 1.1 μm. The 3D reconstruction was performed as previously described. The segmentation of the osteocyte lacunae was achieved with a deep learning-based method using the module available in Dragonfly software in a region of interest of 1.3 mm in thickness centered at the midshaft. 15 slices were used to train the model, followed by a manual refinement.

### Histology

Immediately after dissection, the femurs of 10 different chick embryos were immersed in 70% ethanol and embedded in Polymethylmethacrylate (PMMA, Technovit 9100, Kulzer, Germany) following the protocol of Moreno-Jiméney et al. [49]. Briefly, the mineralized femurs were dehydrated through an ethanol gradient every two days in five steps: 70%, 80%, 96%, 100% and 100%. The samples were then cleared in xylol for 3 h twice at room temperature. The specimens were immersed in a pre-infiltration solution composed of 200 ml of destabilized Technovit 9100 basic solution with 1 g of Hardener 1 for 2 days followed by the infiltration solution (250 ml of destabilized Technovit 9100 basic solution mixed with 20 g of PMMA powder) for 6 days, changing the solution once after 3 days. The final embedding solution was prepared with a 9:1 mixture of a stock solution A (500 ml of destabilized basic solution mixed with 80 g of PMMA powder and 3 g of Hardener 1) and a stock solution B (44 ml of destabilized basic solution with 4 ml of Hardener 2 and 2 ml of regulator). The polymerization occurred for 3 days at – 20°C. The resin blocks were processed for ultrathin sectioning on a Leica microtome (Leica Biosystems Nussloch GmbH, Nussloch, Germany). The histological sections were stained with the Hematoxylin and Eosin rapid kit (Clin-Tech Ltd, Guilford, UK) following the manufacturer guidelines. All the sections were automatically scanned with the mosaic imaging mode using the Keyence VHX-5000 digital microscope (Keyence Corp., Osaka).

### Scanning Electron Microscopy (SEM)

The embedded samples were imaged with a Quanta 600 FEG environmental scanning electron microscope (FEI, Hillsboro, OR, USA) using a backscatter electron (BSE) detector in low vacuum mode (0.5-1 Torr) at 20 kV, without any prior coating.

### Cryo Focused Ion Beam-Scanning Electron Microscopy (FIB-SEM)

Right femoral cross-sections of approximately 500 μm thickness were cut at midshaft using a razor blade in three different chick embryo at EDD13. The bone sections were immediately sandwiched individually between two type B gold-coated copper freezer hats (BALTIC preparation, Wetter, Germany) with the addition of 10 wt% dextran (Sigma, 31390) as cryo-protectant. The freezer hats were positioned in a mirror combination to allow a total cavity thickness of 0.6 mm. The sandwiched samples were cryo-immobilized in a HPM100 high pressure freezing machine (Leica Microsystems, Vienna, Austria) within 5 min after the time of death. The frozen sample carriers were mounted on a cryo sample holder in the Leica EM VCM loading station (Leica Microsystems, Vienna, Austria) at liquid nitrogen temperature. The sample holder was then transferred using the VCT500 shuttle to the ACE600 (Leica Microsystems, Vienna, Austria) for freeze fracture and sputter coating. Subsequently to the freeze-fracture, the exposed samples were sputter coated with an 8 nm platinum layer. Finally, the samples were transferred to the Zeiss Crossbeam 540 (Zeiss Microscopy GmbH, Oberkochen, Germany) using the VCT500 shuttle. Throughout the analysis, the bone samples were kept at a temperature below −145°C.

### Serial Surface View (SSV) imaging

FIB-SEM Serial Surface View Imaging was performed using a Zeiss Crossbeam 540 (Zeiss Microscopy GmbH, Oberkochen, Germany) dual beam microscope (FIB-SEM) in three different femoral midshaft sections. The samples were elevated to the height of 5.1 mm, which corresponds to the coincident point of the two beams and tilted to 54°. A trench of approximately 45 μm length and 75 μm width was milled at 15 nA. The exposed surface was polished and imaged with a lower ion beam current (1.5 nA). The electron beam was then focused on the polished exposed tissue at 2 keV, 50 pA for all stacks. Images were sequentially collected using the “slice and view” protocol after setting the pixel size of the image prior to data collection at 8 nm in the x and y directions. The slice thickness (z direction) was also set at 8 nm to maintain an isometric voxel size. All stacks were acquired with an image resolution of 4096 x 3072 pixels, in an 8-bit greyscale images and using the dual channel option. This option allows the simultaneous acquisition of images from mixed Inlens/secondary electron (SE) detector signal and the backscattered electron (BSE) detector. Stack 1 comprises 1625 slices; stack 2, 1876 slices; and stack 3, 303 slices.

### FIB-SEM image processing and segmentation

Stacks alignment, image processing, and segmentations were generated using Dragonfly software, Version 2021.1 (Object Research Systems (ORS) Inc, Montreal, Canada). The videos were first generated in Dragonfly software and then edited in Movavi Video Editor 15 Plus software (Movavi, Saint Louis, MO, USA). Both BSE and mixed Inlens/SE images were first automatically aligned using the sum of square differences (SSD) method available in the slice registration panel. Curtaining artifacts produced by the beam during the milling process were corrected using the vertical destriping filter. To improve the visualization of the structure, the contrast was enhanced, and the noise reduced using a convolution filter in all stacks. Contrast was further improved in the mixed Inlens/SE images of all stacks using the contrast limited adaptive histogram equalization (CLAHE) filter.

The segmentation of the denser, brighter parts as well as the canaliculi was performed in the BSE stacks with the deep learning module in Dragonfly software (Object Research Systems (ORS) Inc, Montreal, Canada) using 20 slices for training each data set, followed by a manual refinement using the ‘brush’ tool. In the mixed SE/Inlens stacks, the deep learning module was also employed to segment the secondary channels and the vesicles, the nucleus and the cells were segmented using the active contour method available in Dragonfly software (Object Research Systems (ORS) Inc, Montreal, Canada).

## Acknowledgements

We thank Daniel Werner for technical support with the micro-CT imaging, Jeannette Steffen for histology staining and Birgit Schonert for sample polishing (all in the Department of Biomaterials at the Max Planck Institute of Colloids and Interfaces). PF and RW acknowledge support from the Max Planck Queensland Centre for the Materials Science of Extracellular Matrices.

