## Supporting Information for "Logistics of Bone Mineralization in the Chick Embryo Studied by 3D Cryo FIB-SEM Imaging"

Movie legends:

**Movie S1:** Perspective rendering of the images acquired in stack 2. The movie shows first, the processed mixed Inlens/SE images and then the BSE images. The stack highlights the 3D arrangement of the mineralized bone, the vesicles containing mineral precursors and the bone cells based on the segmentation of these structures. Pause 1: Enlarged processed image of the mixed Inlens/SE detectors taken from the acquisition plan (xy) showing the bone and the organic matrix. A nucleus is clearly identifiable. Pause 2: Enlarged processed image of the mixed Inlens/SE detectors taken from the reconstruction plan (yz) showing the bone and the organic matrix in a quality similar to the acquisition plan due to the isometric and high-resolution voxel size (8nm). Pause 3: Enlarged processed image of first the BSE detector showing the mineralized bone (yellow), the rough contour of the cells (dotted orange) and their nucleus when present (turquoise). The green arrows point to mineral precursors that appear brighter in the BSE images. The same image from the mixed Inlens/SE detector shows that the mineral precursors identified in the BSE image are found within numerous vesicles and can be identified with a granular structure. Unmineralized collagen fibrils can be seen between the cells and the mineralized bone.

**Movie S2:** The movie shows the same stack as in Movie 1 with the segmented data. The nanochannels (blue) form an interconnected network with the canaliculi (red) in the mineralized bone matrix (light yellow). Numerous vesicles (purple) containing mineral precursors (green) are found inside the cells (orange) close to the mineralized bone matrix.

**Movie S3:** Perspective rendering of a complete femur at EDD13, contrast-stained with phosphotungstic acid. The movie shows then the segmentation of the osteocyte lacunae performed on a region of interest of 1.3 mm in thickness, corresponding to a bone volume of 0.2028 $mm^3$. A total of 39 716 osteocytes lacunae were counted, which gives a density of 196 000 osteocytes lacunae/ $mm^3$.

| **FIB SEM stack 1** | | | | | | | | |
|---|---|---|---|---|---|---|---|---|
| Volume imaged | 8nm voxel | 4096*3072 store resolution | 1625 images | 32.77 µm x24.58µm x13 µm | 10 471.33 $\mu m^3$ | | | |
| **Segmentation** | | | | | | | | |
| Mineralized bone | 2734.51 $\mu m^3$ | | | | | | | |
| Cells Total | 4693.00 $\mu m^3$ | Number of cells: 6 | | | | | | |
| Canaliculi | 59.27 $\mu m^3$ | | | | | | | |
| Nanochannels channels | 398.11$\mu m^3$ | | | | | | | |
| Total vesicles | 72.71$\mu m^3$ | Number of vesicles: 122 | Minimum: 0.01 $\mu m^3$ | Maximum: 3.47 $\mu m^3$ | SD: 0.55 $\mu m^3$ | Mean: 0.60 $\mu m^3$ | | |
| Total mineral precursors | 7.11$\mu m^3$ | Minimum: 0.001$\mu m^3$ | Maximum: 0.45$\mu m^3$ | SD: 0.07$\mu m^3$ | Mean: 0.06$\mu m^3$ | | | |
| Cell 1: | 304.15$\mu m^3$ | Volume Nucleus: 13.35 $\mu m^3$ | Number of vesicles: 5 | Total volume of vesicles: 2.75 $\mu m^3$ | Minimum: 0.2 $\mu m^3$ | Maximum: 1.08 $\mu m^3$ | SD: 0.44 $\mu m^3$ | Mean: 0.55 $\mu m^3$ |
| | | | | Total mineralization: 0.10 $\mu m^3$ | Minimum: 0.002 $\mu m^3$ | Maximum: 0.05 $\mu m^3$ | SD: 0.02 $\mu m^3$ | Mean: 0.02 $\mu m^3$ |
| Cell 2: | 296.95$\mu m^3$ | Volume Nucleus: 9.15 $\mu m^3$ | Number of vesicles: 6 | Total volume of vesicles: 10.20 $\mu m^3$ | Minimum: 0.66 $\mu m^3$ | Maximum: 3.47 $\mu m^3$ | SD:0.88 $\mu m^3$ | Mean: 1.70 $\mu m^3$ |
| | | | | Total mineral precursors: 1.13 $\mu m^3$ | Minimum: 0.06 $\mu m^3$ | Maximum: 0.45 $\mu m^3$ | SD: 0.13 $\mu m^3$ | Mean: 0.19 $\mu m^3$ |
| Cell 3: | 469.13 $\mu m^3$ | Volume Nucleus: 18.53 $\mu m^3$ | No vesicles | | | | | |
| Cell 4: | 212.46 $\mu m^3$ | No nucleus | No vesicles | | | | | |
| Cell 5: | 1445.69 $\mu m^3$ | No nucleus | Number of vesicles: 32 | Total volume of vesicles: | Minimum: 0.13 $\mu m^3$ | Maximum: 1.69 $\mu m^3$ | SD: 0.36 $\mu m^3$ | Mean: 0.58 $\mu m^3$ |

| | | | | | | | | |
|---|---|---|---|---|---|---|---|---|
| | | | | 17.91 µm$^3$ | | | | |
| | | | | Total mineral precursors: 1.84 µm$^3$ | Minimum: 0.006 µm$^3$ | Maximum: 0.33 µm$^3$ | SD: 0.06 µm$^3$ | Mean: 0.06 µm$^3$ |
| Cell 6: | 1905.67 µm$^3$ | Volume Nucleus: 43.21 µm$^3$ | Number of vesicles: 79 | Total volume of vesicles: 41.86 µm$^3$ | Minimum: 0.02 µm$^3$ | Maximum: 2.70 µm$^3$ | SD: 0.49 µm$^3$ | Mean: 0.53 µm$^3$ |
| | | | | Total mineral precursors: 4.03 µm$^3$ | Minimum: 0.001 µm$^3$ | Maximum: 0.32 µm$^3$ | SD: 0.07 µm$^3$ | Mean: 0.07 µm$^3$ |
| **FIB SEM stack 2** | | | | | | | | |
| Volume imaged | 8nm voxel | 4096*3072 store resolution | 1876 images | 32.768µm x 24.576µm x 15.008µm | 12 086.038 µm$^3$ | | | |
| **Segmentation** | | | | | | | | |
| Mineralized bone | 2626.39 µm$^3$ | | | | | | | |
| Cells Total | 8164.10 µm$^3$ | Number of cells: 6 | | | | | | |
| Canaliculi | 88.23 µm$^3$ | | | | | | | |
| Nanochannels channels | 378.27 µm$^3$ | | | | | | | |
| Total vesicles | 199.14 µm$^3$ | Number of vesicles: 270 | Minimum: 0.03 µm$^3$ | Maximum: 5.14 µm$^3$ | SD: 0.66 µm$^3$ | Mean: 0.74 µm$^3$ | | |
| Total mineral precursors | 16.29 µm$^3$ | Minimum: 0.0004 µm$^3$ | Maximum: 0.64 µm$^3$ | SD: 0.07 µm$^3$ | Mean: 0.06 µm$^3$ | | | |
| Cell 1: | 359.47 µm$^3$ | Volume Nucleus: 181.49 µm$^3$ | Number of vesicles: 2 | Total volume of vesicles: 2.17 µm$^3$ | Minimum: 1.06 µm$^3$ | Maximum: 1.11 µm$^3$ | SD: 0.03 µm$^3$ | Mean: 1.09 µm$^3$ |
| | | | | Total mineral precursors: 0.08 µm$^3$ | Minimum: 0.004 µm$^3$ | Maximum: 0.07 µm$^3$ | SD: 0.03 µm$^3$ | Mean: 0.04 µm$^3$ |
| Cell 2: | 822.78 µm$^3$ | No nucleus | Number of vesicles: 14 | Total volume of vesicles: 8.02 µm$^3$ | Minimum: 0.09 µm$^3$ | Maximum: 1.58 µm$^3$ | SD: 0.46 µm$^3$ | Mean: 0.57 µm$^3$ |

| | | | | | | | | |
|---|---|---|---|---|---|---|---|---|
| | | | | Total mineral precursors: 0.67 µm$^3$ | Minimum: 0.03 µm$^3$ | Maximum: 0.18 µm$^3$ | SD: 0.06 µm$^3$ | Mean: 0.06 µm$^3$ |
| Cell 3: | 1221.70 µm$^3$ | Volume Nucleus: 57.91 µm$^3$ | Number of vesicles: 29 | Total volume of vesicles: 37.40 µm$^3$ | Minimum: 0.05 µm$^3$ | Maximum: 5.14 µm$^3$ | SD: 1.19 µm$^3$ | Mean: 1.29 µm$^3$ |
| | | | | Total mineral precursors: 1.46 µm$^3$ | Minimum: 0.01 µm$^3$ | Maximum: 0.22 µm$^3$ | SD: 0.06 µm$^3$ | Mean: 0.05 µm$^3$ |
| Cell 4: | 134.48 µm$^3$ | No nucleus | Number of vesicles: 1 | Total volume of vesicles: 1.40 µm$^3$ | | | | |
| | | | | Total mineral precursors: 0.01 µm$^3$ | | | | |
| Cell 5: | 3564.34 µm$^3$ | Volume Nucleus: 119.49 µm$^3$ | Number of vesicles: 146 | Total volume of vesicles: 106.30 µm$^3$ | Minimum: 0.03 µm$^3$ | Maximum: 3.25 µm$^3$ | SD: 0.57 µm$^3$ | Mean: 0.73 µm$^3$ |
| | | | | Total mineral precursors: 9.95 µm$^3$ | Minimum: 0.0004 µm$^3$ | Maximum: 0.64 µm$^3$ | SD: 0.08 µm$^3$ | Mean: 0.08 µm$^3$ |
| Cell 6: | 2064.36 µm$^3$ | Volume Nucleus: 119.08 µm$^3$ | Number of vesicles: 78 | Total volume of vesicles: 43.85 µm$^3$ | Minimum: 0.07 µm$^3$ | Maximum: 2.90 µm$^3$ | SD: 0.41 µm$^3$ | Mean: 0.56 µm$^3$ |
| | | | | Total mineral precursors: 4.10 µm$^3$ | Minimum: 0.0004 µm$^3$ | Maximum: 0.29 µm$^3$ | SD: 0.06 µm$^3$ | Mean: 0.06 µm$^3$ |
| **FIB SEM stack 3** | | | | | | | | |
| Volume imaged | 8nm voxel | 4096*2460 store resolution | 303 images | 32.77 µm x19.68 µm x2.424 µm | 1563.3 µm$^3$ | | | |
| **Segmentation** | | | | | | | | |
| Mineralized bone | 767.87 µm$^3$ | | | | | | | |

| | | | | | | | | |
|---|---|---|---|---|---|---|---|---|
| Cells Total | 473.49 $\mu m^3$ | Number of cells: 7 | | | | | | |
| Canaliculi | 7.93 $\mu m^3$ | | | | | | | |
| Nanochannels channels | 109.39 $\mu m^3$ | | | | | | | |
| Total vesicles | 10.20 $\mu m^3$ | Number of vesicles: 40 | Minimum: 0.002 $\mu m^3$ | Maximum: 1.04 $\mu m^3$ | SD: 0.25 $\mu m^3$ | Mean: 0.24 $\mu m^3$ | | |
| Total mineral precursors | 1.01 $\mu m^3$ | Minimum: 0.001 $\mu m^3$ | Maximum: 0.21 $\mu m^3$ | SD: 0.004 $\mu m^3$ | Mean: 0.02 $\mu m^3$ | | | |
| Osteocyte 1: | 114.44 $\mu m^3$ | Volume Nucleus: 48.24 $\mu m^3$ | Number of vesicles: 13 | Total volume of vesicles: 3.25 $\mu m^3$ | Minimum: 0.04 $\mu m^3$ | Maximum: 0.68 $\mu m^3$ | SD: 0.18 $\mu m^3$ | Mean: 0.27 $\mu m^3$ |
| | | | | Total mineral precursors: 0.25 $\mu m^3$ | Minimum: 0.002 $\mu m^3$ | Maximum: 0.05 $\mu m^3$ | SD: 0.01 $\mu m^3$ | Mean: 0.02 $\mu m^3$ |
| Osteocyte 2: | 86.72 $\mu m^3$ | Volume Nucleus: 29.37 $\mu m^3$ | Number of vesicles: 9 | Total volume of vesicles: 2.25 $\mu m^3$ | Minimum: 0.05 $\mu m^3$ | Maximum: 1.04 $\mu m^3$ | SD: 0.37 $\mu m^3$ | Mean: 0.34 $\mu m^3$ |
| | | | | Total mineral precursors: 0.20 $\mu m^3$ | Minimum: 0.001 $\mu m^3$ | Maximum: 0.10 $\mu m^3$ | SD: 0.03 $\mu m^3$ | Mean: 0.03 $\mu m^3$ |
| Osteocyte 3: | 27.53 $\mu m^3$ | No nucleus imaged | No vesicles inside | | | | | |
| Cell 4: | 111.04 $\mu m^3$ | Volume Nucleus: 68.18 $\mu m^3$ | Number of vesicles: 3 | Total volume of vesicles: 1.27 $\mu m^3$ | Minimum: 0.19 $\mu m^3$ | Maximum: 0.84 $\mu m^3$ | SD: 0.30 $\mu m^3$ | Mean: 0.42 $\mu m^3$ |
| | | | | Total mineral precursors: 0.02 $\mu m^3$ | Minimum: 0.01 $\mu m^3$ | Maximum: 0.01 $\mu m^3$ | SD: 0.001 $\mu m^3$ | Mean: 0.01 $\mu m^3$ |
| Cell 5: | 45.79 $\mu m^3$ | Volume Nucleus: 9.83 $\mu m^3$ | Number of vesicles: 7 | Total volume of vesicles: 2.48 $\mu m^3$ | Minimum: 0.06 $\mu m^3$ | Maximum: 0.91 $\mu m^3$ | SD: 0.30 $\mu m^3$ | Mean: 0.35 $\mu m^3$ |

| | | | | | | | | |
|---|---|---|---|---|---|---|---|---|
| | | | | Total mineral precursors: 0.44 $\mu m^3$ | Minimum: 0.01 $\mu m^3$ | Maximum: 0.21 $\mu m^3$ | SD: 0.06 $\mu m^3$ | Mean: 0.06 $\mu m^3$ |
| Cell 6: | 66.19 $\mu m^3$ | No nucleus imaged | Number of vesicles: 6 | Total volume of vesicles: 0.8 $\mu m^3$ | Minimum: 0.01 $\mu m^3$ | Maximum: 0.40 $\mu m^3$ | SD: 0.12 $\mu m^3$ | Mean: 0.15 $\mu m^3$ |
| | | | | Total mineral precursors: 0.07 $\mu m^3$ | Minimum: 0.001 $\mu m^3$ | Maximum: 0.03 $\mu m^3$ | SD: 0.01 $\mu m^3$ | Mean: 0.01 $\mu m^3$ |
| Cell 7: | 21.78 $\mu m^3$ | No nucleus imaged | Number of vesicles: 2 | Total volume of vesicles: 0.13 $\mu m^3$ | Minimum: 0.06 $\mu m^3$ | Maximum: 0.08 $\mu m^3$ | SD: 0.01 $\mu m^3$ | Mean: 0.07 $\mu m^3$ |
| | | | | Total mineral precursors: 0.02 $\mu m^3$ | Minimum: 0.01 $\mu m^3$ | Maximum: 0.01 $\mu m^3$ | SD: 0.003 $\mu m^3$ | Mean: 0.01 $\mu m^3$ |

**Table S1:** Raw quantitative data for all three stacks investigated. All volumes and counts are based on the segmentation of the different structures using Dragonfly software (Object Research Systems (ORS) Inc, Montreal, Canada).

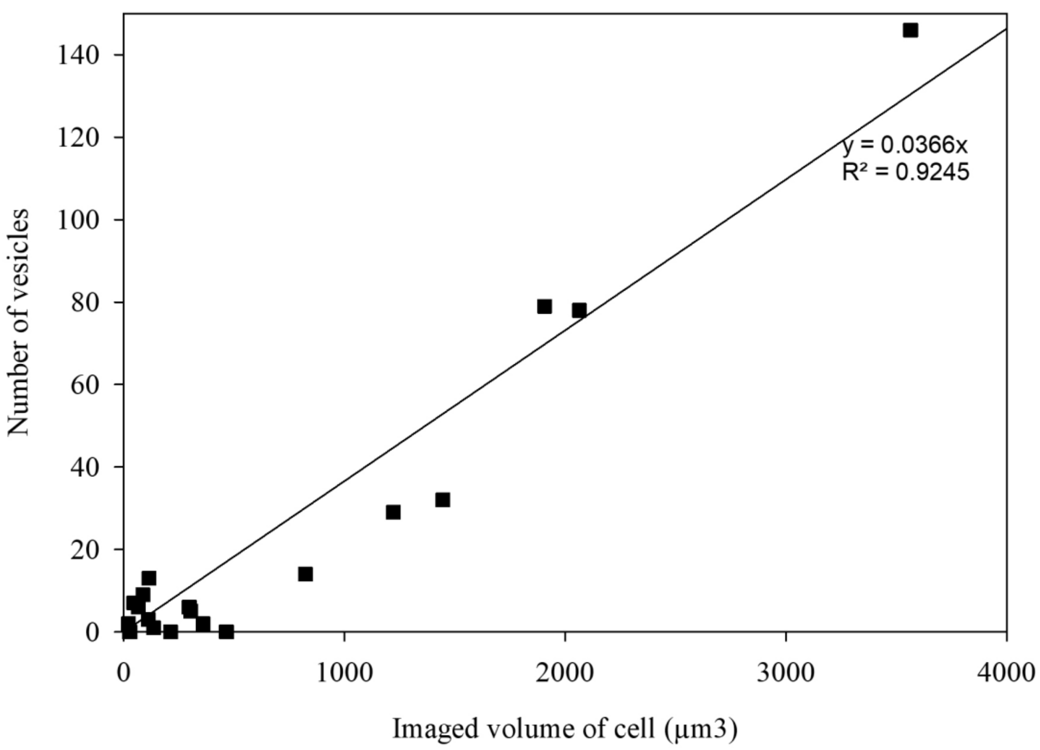


**Figure S1:** Linear regression of the number of vesicles according to the volume of cell partially imaged for the three stack combined. The pre-factor of 0.0366 correspond to the vesicles density.
